# Embryological insights into the evolution of genome regulation using haploid and diploid whiteflies, *Bemisia tabaci*

**DOI:** 10.1101/2024.07.30.605827

**Authors:** Emily A. Shelby, Elizabeth C. McKinney, Alvin M. Simmons, Allen J. Moore, Patricia J. Moore

**Affiliations:** Department of Entomology, University of Georgia, 120 Cedar Street, Athens, Georgia, 30602, USA; U.S. Department of Agriculture, Agricultural Research Service, U.S. Vegetable Laboratory, Charleston, South Carolina, 29414, USA

**Keywords:** *DNA methyltransferase 1*, Embryogenesis, Haplodiploid, *Bemisia tabaci*, RNA interference

## Abstract

The whitefly, *Bemisia tabaci*, is a hemipteran with a haplodiploid sex determination system, which provides a natural experiment for examining genome function in haploid and diploid embryos. Yet the embryogenesis of *B. tabaci* remains understudied. Our previous work has established a possible role of *DNA methyltransferase 1* (*Dnmt1*) in genome stability. In this study we used maternal RNA interference (RNAi) and immunohistochemistry to study the complex dynamics between DNMT1 and ploidy during embryogenesis. We found that both haploid and diploid *B. tabaci* have a similar developmental timeline to other holometabolous insects. We also found that, like other obligatory haploid insects, ploidy does not affect developmental timing, suggesting that maternal factors play a greater role than ploidy in developmental timing. For embryos with reduced expression of *Dnmt1*, we found that the loss of DNMT1 disrupts blastoderm development and affects nuclei morphology in both haploid and diploid embryos. Our results suggest that DNMT1 is required for blastoderm development.

## 1. Introduction

The whitefly, *Bemisia tabaci*, is a globally-important pest, causing billions of US dollars in damage to crops each year (Stansly and Naranjo, 2010, Inoue-Nagata et al., 2016, Czosnek et al., 2017). As such, considerable work has been directed towards their ecological impacts and possible pest-management strategies (Shelby et al., 2020). *Bemisia tabaci* is difficult to manage using traditional chemical means such as neonicotinoid pesticides (Horowitz et al., 2020). This leads to a critical need to ascertain the basic biological functions in this insect that will, in turn, facilitate effective management strategies. Deciphering developmental and molecular mechanisms underlying the life history of *B. tabaci* can be useful in this regard and allow researchers to optimize applications of modern molecular technologies for use in pest management (Shelby et al., 2020; Suhag et al., 2020). Indeed, the importance of *B. tabaci* as a pest has led to an increase in the number of available technologies that make whitefly molecular studies more tractable such as a sequenced genomes (Chen at al., 2016; Xie et al., 2017; Chen et al., 2019; Hussain et al., 2019), a characterized methylome (Cunningham et al., 2024), RNA sequencing data sets (Shen et al., 2023; Cunningham et al., 2024;), transcriptomic and proteomic data sets (Yang et al., 2013), protocols for CRISPR (Heu et al., 2020) and RNA interference (RNAi; Dai et al., 2017; Gong et al., 2022; Jain et al., 2022; Shelby et al., 2023), life history summaries (Byrne et al., 1991; Aregbesola et al., 2020), and imaging methods (Luan et al., 2018; Bondy and Hunter, 2019; Shelby et al., 2023). The unique biology of this whitefly also lends itself to asking fundamental questions in novel yet widely-applicable ways. For example, *B. tabaci*, as a member of the order Hemiptera, is assumed to undergo embryo development with in a similar time frame and using the same embryo patterning scheme. However, *B. tabaci* uses a haplodiploid sex determination system, which is almost exclusively studied in Holometabola and is known to affect developmental patterning and morphogenesis (Netschitailo et al., 2022). However, due to a lack of embryological studies, it is unknown to what extent hemimetabolous embryo development is evolutionarily conserved in *B. tabaci* or how their sex determination system affects their patterns of development.

Nucleocytoplasmic (N/C) ratio, or the amount of cytoplasm relative to the DNA content, influences timing of the early embryonic events such as cellularization and changes in the cell cycle length that result in zygotic genome activation (Schier, 2007). In haplodiploid systems unfertilized eggs become haploid males (n) and fertilized eggs become diploid females (2n). Thus, one question is how is developmental timing is regulated in the same species with differences in ploidy? In *Drosophila* embryos where development is artificially activated and ploidy can be manipulated, haploid embryo development is slower and the maternal-to-zygotic transition (MZT) associated with cellularization takes longer to occur (Edgar et al., 1986; Erickson and Quintero, 2007; Shermoen, McCleland, and O’Farrell, 2010). However, in Hymenoptera such as *Nasonia*, where a haplodiploid sex determination system is typical and normal, ploidy does not affect when embryonic events such as cellularization and zygotic genome activation occur, suggesting that maternal factors determining the timing of these events (Arsala and Lynch, 2017). It is uncertain to what extent ploidy or maternal factors regulate *B. tabaci* early embryo development.

Previous work by our laboratory suggests that one fruitful avenue for investigating factors related to both insect embryogenesis and genome regulation is *DNA methyltransferase I (Dnmt1)* function. *Dnmt1* is characterized as a maintenance methyltransferase based on its affinity for hemi-methylated DNA (Goll and Bestor, 2005).

However, functional manipulation suggests DNA methylation is not the essential mechanism through which DNMT1 is acting in insects (Schulz et al., 2018; Amukamara et al., 2020; Ivasyk et al., 2023; Shelby et al., 2023). A methylation-independent role is not improbable. It has been hypothesized in mammal systems that *Dnmt1* plays a role in DNA repair and chromosome stability during the cell cycle in addition to its canonical role as a methyltransferase (Brown & Robertson, 2007). While there is mounting evidence that *Dnmt1* has a conserved role in these processes (Mortusewicz et al., 2005; Loughery et al., 2011; Uysal et al., 2015;Madakashira et al., 2024), the methylation-dependent and methylation-independent functions cannot be separated because DNA methylation is necessary for mammalian gene expression (Li & Zhang, 2014). Insects, therefore, offer a unique window into possible non-canonical functions of DNMT1 in a taxa that does not rely heavily on DNA methylation.

Although the importance of DNA methylation in insects is unclear (Bewick et al. 2017, Duncan et al., 2022), DNMT1, the enzyme associated with maintaining DNA methylation, is vital for proper gametogenesis and embryogenesis (Zwier et al., 2011; Kay et al., 2018; Schulz et al., 2018; Bewick et al., 2019; Amukamara et al., 2020; Ventós-Alfonso, 2020; Washington et al., 2021; Arsala et al., 2022; Ivasyk et al., 2023 Shelby et al., 2023). Due to a lack of studies that look beyond the superficial loss of reproductive capabilities, the mechanisms behind DNMT1’s seemingly methylation-independent function in insect embryogenesis is unknown. In *B. tabaci*, knockdown of *Dnmt1* results in the production of fewer eggs as well as fewer eggs successfully hatching (Shelby et al., 2023). Though there was evidence that these eggs were abnormal prior to oviposition (Shelby et al., 2023), it is unclear whether the eggs failed to develop because they lacked maternal factors, and embryonic development could never occur in the first place, or if embryonic development began to occur but was prohibited from successfully progressing. It is also unclear what the role of ploidy was in this knockdown phenotype because all embryos in Shelby et al. (2023) were haploid.

The aim of our study was to investigate mechanisms that affect genome function during embryogenesis in *B. tabaci*. We approached this by looking at the complex relationship between ploidy and *Dnmt1* in the context of early insect embryonic development. First, we chronicled embryonic development in order to get a detailed *B. tabaci* developmental timeline. Then we performed maternal RNAi followed by mating experiments to produce either “haploid-only” or “haploid or diploid” embryos. Finally, we used immunohistochemistry to compare differences in early embryonic development, including differences in nuclei size, in the context of *Dnmt1*-knockdown.

## 2. Materials and Methods

### 2.1. Animal husbandry

All individuals used in the experiments were taken from laboratory reared MEAM1 *B. tabaci* colonies. The colonies were started from wild caught populations collected from cotton fields in Tift County, Georgia, USA in 2018 (McKenzie et al., 2020). In the laboratory, we reared the colonies on collard plants (*Brassica oleracea*). We kept both *B. tabaci* and collards in an incubator at 27ºC with a 14:10 hour light: dark photoperiod and constant temperature and photoperiod throughout experiments. We removed newly-eclosed females from nymph colonies within 24 hours of eclosion to verify female age and mating status. For experiments requiring untreated females (i.e. females not fed dsRNA), we placed unmated females 3-5 days post adult eclosion either on plants in a female-only colony (to create “haploid-only” embryos) or on plants with males (to create “haploid or diploid” embryos). Diploid embryos are the product of sexual reproduction from mating. However, not every offspring produced after mating will be diploid. In our colony, approximately 38.9% of all offspring produced from mated females developed into diploid females. Therefore, embryos from mated females are classified as “haploid or diploid.” For females designated for treatment, the newly eclosed females were put in female-only colonies where they remained until receiving dsRNA treatment.

### 2.2. Maternal RNA interference (RNAi)

We used the same double stranded interfering RNA and knockdown protocol as that in Shelby et al. 2023. Briefly, we made DNA templates of *Dnmt1* and *eGFP* using PCR amplification with gene-specific primers and 500 ng/µl RNA (Table 1). We synthesized sense and antisense RNA in a single reaction using the Ambion MEGAscript RNAi kit (ThermoFischer Sci, Waltham, MA, USA) following the manufacturer’s instructions. Following extraction and ethanol precipitation, we aliquoted the double-stranded RNA (dsRNA) and stored it at -80ºC until use.

**Table 1.**
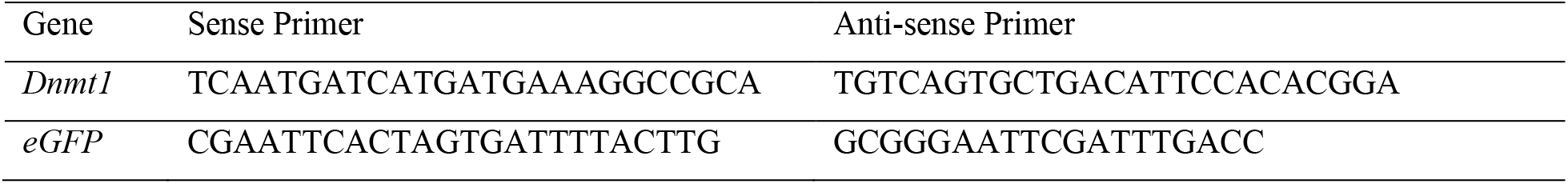
Primer sequences used for dsRNA synthesis.

For both “haploid-only” and “haploid or diploid” groups, we used virgin females 3-5 days post adult eclosion. We then treated females with a dsRNA solution using an artificial feeding mechanism previously described in Shelby et al. (2023). The dsRNA feeding solution contained green food coloring (McCormick & Company, Baltimore, MD, USA). The food coloring allowed us to confirm if a female had fed on the solution. If the females did feed on the solution, their abdomens would appear green when observed under a microscope. Only females we confirmed had imbibed the dsRNA solution were used for experiments.

Following feeding, we placed the “haploid-only” producing females on 15 cm tall collard plants. Each plant harbored 50-100 treated females. We checked plants for eggs every six hours. If eggs were present, we removed females from the plant and placed them on a new plant. We placed “haploid or diploid” producing females on 15cm tall collard plants with untreated males 3-5 days post adult eclosion. Each plant harbored 50-100 treated females along with an equal number of untreated males. We checked plants for eggs every six hours. If eggs were present, we removed females and males were removed from the current plant and placed them on a new plant.

### 2.3. Immunohistochemistry and image acquisition

Following oviposition, we collected collard leaves with eggs. We either immediately removed the eggs from the leaves or kept them on the leaves until they were the appropriate age. In the case of eggs kept on leaves, we wrapped the leaves in damp paper towel and placed them in a petri dish back in the incubator to prevent desiccation. For our embryonic development overview study, we used eggs at approximately 6, 12, 18, 24, 48, 72, and 84 hours post oviposition (PO). For ploidy and knockdown studies, we used eggs at approximately 6, 12, 18, and 24 hours PO. Eggs were kept separate from adults.

We removed eggs from the leaves using a probe mounted with a minute pin. After removal from the leaves, we dechorionated eggs by soaking them in 5% sodium hypochlorite for 15 minutes. Following the removal of the chorion, we washed embryos in PBS and incubated in 32% paraformaldehyde overnight at 2°C. To visualize nucleic acids, we used 1 μL 0.5 μg/mL, DAPI in PBS (Thermofisher Scientific, Waltham, MA, USA). We mounted the stained embryos with Mowiol 4-88 mounting medium (Sigma-Aldrich, St. Louis, MO, USA).

We imaged the embryos with a Zeiss LSM 880 Confocal Microscope (Zeiss) at the University of Georgia Biomedical Microscope Core. Z-stack maximum projection images were taken. Only global image adjustment features (such as brightness and contrast) were used. All confocal images were falsely colored.

### 2.4. Embryo staging

Embryogenesis in *B. tabaci* has not been previously characterized. Therefore, we established a staging system to morphologically and chronologically identify key developmental stages from the initial syncytial cleavage to completion of the embryo body plan. We divided early embryogenesis into five main stages based morphological hallmarks described in other hemipterans. These stages were: cleavage, blastoderm, gastrulation, segmentation, and growth and later development. The timing of each stage is presented as a range based on how many hours PO the stage is observed. To be consistent with previous experiments (Shelby et al., 2023), we developed this developmental timeline using eggs from untreated virgin females, which produce “haploid-only” embryos. Anterior-posterior polarity of the egg is easily identified due to the large bacteriocyte located at the posterior portion of the egg. Bacteriocyte position is established during oogenesis in the mature oocytes (Shelby et al., 2023).

### 2.5 Image analysis

We successfully processed and analyzed a total of N = 149 embryos, which included N = 100 embryos from wildtype females, N = 14 embryos from “haploid-only” *dseGFP*-fed females, N = 10 embryos from “haploid-only” *dsDnmt1*-fed females, N = 15 embryos from “haploid or diploid” *dseGFP-*fed females, and N = 10 embryos from “haploid or diploid” *dsDnmt1*-fed mated females.

For nuclei measurements, we used the ImageJ measuring software (Version 1.54i by FIJI). For embryos with less than 30 nuclei, we measured all nuclei. For embryos with more than 30 nuclei, we measured 30 randomly-selected nuclei.

### 2.6. Statistical analyses

We used a one-way ANOVA to compare the variation in the size of embryo nuclei. For data analyses, we used Base R in RStudio (RStudio Team, 2023). We used the ggplot package for data visualization (Wickman, 2016).

## 3. Results

### 3.1. Overview of B. tabaci embryonic development

Like other hemipterans, *B. tabaci* embryos develop as a syncytium during the cleavage stage from 0-12 hours post-oviposition (PO). Syncytial cleavage begins initially at the center of the egg around (Fig. 1A) and expands outward towards the periphery (Fig. 1B). During this time the nuclei divide synchronously, and the size of the nuclei vary as they rapidly progress through the cell cycle (as seen in Fig. 1A-C). After reaching the periphery, the nuclei are regularly-spaced and are approximately the same size, suggesting that they are undifferentiated (Fig. 1D).

**Fig. 1.**
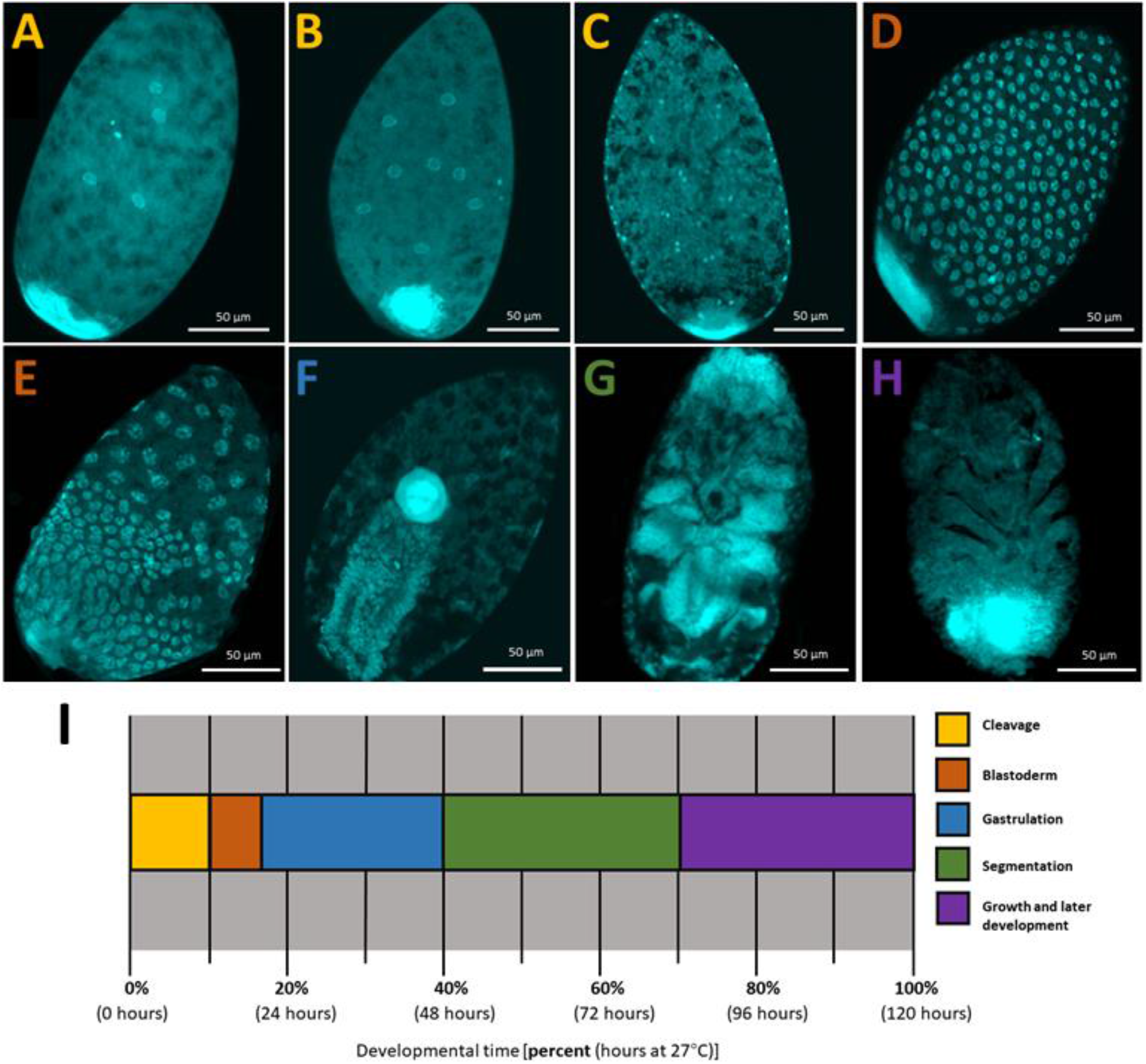
Illustration of key developmental stages during *B. tabaci* embryogenesis by DAPI nuclear staining: (A-C) Cleavage, (D-E) Blastoderm, (F) Gastrulation, (G) Segmentation, and (H) Growth and later development. (I) A developmental timeline illustrating the onset and duration of stages examined in this study (color of letter denoting embryo image corresponds with the stage color on the timeline).

During the blastoderm stage (12-18 hours PO) membranes begin to form around the nuclei. Concurrently, two cell populations begin to segregate: blastoderm cells and extraembryonic cells. The blastoderm cells concentrate towards the posterior region near the bacteriocyte, forming the germ rudiment (Fig. 1E).

Gastrulation takes place from 18 hours PO to 48 hours PO. This stage begins with the condensation of the germ rudiment and invagination towards the center (Fig. 1F). The bacteriocyte also moves towards the center. As a result of this movement, the embryo extends with the cephalic region at the posterior region of the egg.

Segmentation occurs approximately 48-84 hours PO. Cephalic and thoracic appendages are visible first (Fig. 1G), followed by abdominal segments housing the bacteriocyte. For most of segmentation, the embryo is immersed in yolk. Subsequent stages of growth and development occur until approximately 120 hours PO when the embryo hatches as a nymph. These stages include elongation of appendages and the dorsal closure of the embryo (Fig. H). During this stage, visualization with fluorescent nuclear staining is disturbed, likely due to secretion of the cuticle.

### 3.2. Effects of ploidy on developmental timing in embryos from wildtype females

To account for how the timing of developmental stages may be affected by ploidy, we compared development between “haploid-only” and “haploid or diploid” embryos. Of the 50 “haploid or diploid” embryos examined from wildtype mated females (N = 10 for each timepoint), developmental timing was consistent with that of “haploid-only” embryos (Table 2). Even if some of the eggs were haploid, there were no obvious outliers suggesting differences between “haploid-only” or “haploid and diploid” embryos. In the “haploid and diploid” embryos, syncytial cleavage was observed at six hours PO. Similarly, cellularization and blastoderm formation occurred by 18 hours PO. Gastrulation had occurred in all embryos by 24 hours PO. Based on our observations, we concluded that ploidy did not have an effect on the timing of embryonic development in embryos from wildtype females.

**Table 2.** Development of “haploid or diploid” embryos from mated wildtype females.

| Hours PO | Phenotype | Stage | n |
| --- | --- | --- | --- |
| 6 | Center-located<br>nuclei | Cleavage | 10 |
| 12 | Nuclei<br>throughout | Cleavage | 9 |
| 18 | Germ rudiment<br>formation | Blastoderm | 8 |
| 24 | Bacteriocyte<br>moves inward | Gastrulation | 10 |
| 48 | Invagination<br>completed | Gastrulation | 10 |

### 3.3. Effects of Dnmt1 on early embryonic development

To investigate the role of *Dnmt1* during embryogenesis, we examined embryo development following manipulation of *Dnmt1* expression by RNAi. Maternal RNAi knockdown of *Dnmt1* produced lethal phenotypes in 70% of embryos. These embryos did not develop beyond the blastoderm stage (18 hours PO) and failed to form a germ rudiment (Fig. 2). The remaining 30% of embryos did not have a knockdown phenotype; this is consistent with egg viability measurements previously reported in Shelby et al. (2023). At 24 hours PO, both “haploid-only” and “haploid or diploid” embryos from *dsDnmt1*-fed females had the same knockdown phenotype (Fig. 2D & L). In contrast, “haploid-only” and “haploid or diploid” embryos from the control *dseGFP*-fed females exhibited a phenotype indistinguishable from embryos produced by wildtype female and proceeded to undergo gastrulation at 24 hours PO (Fig. 2H& P). These results suggest that *Dnmt1* may be required for cell differentiation and germ rudiment formation.

**Fig. 2.**
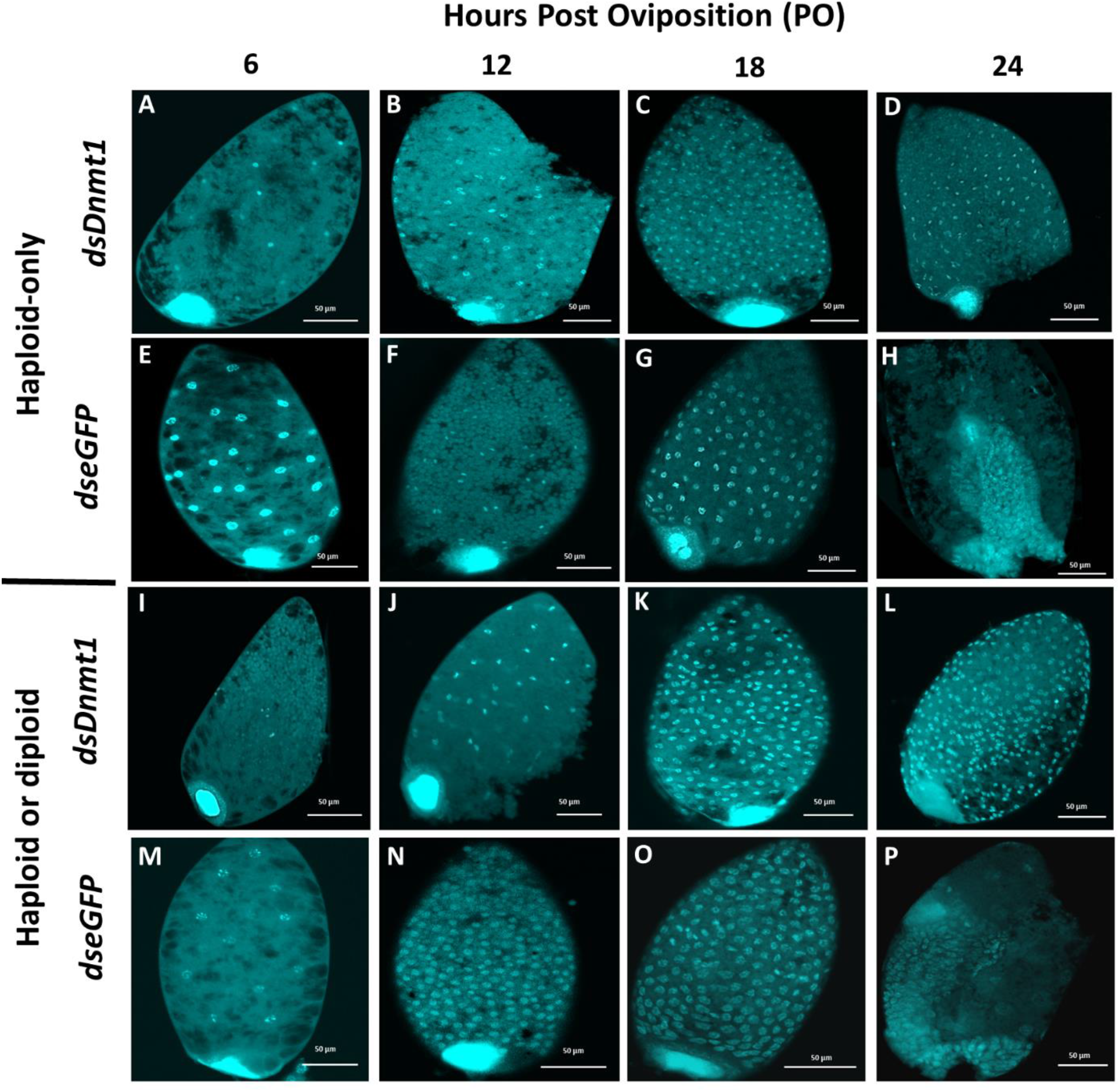
Knockdown of *Dnmt1* via maternal RNAi results in inability to complete cellularization and form a germ rudiment at 24 hours post oviposition (PO). Images were taken at 6, 12, 18, and 24 hours PO of (A-D) *dsDnmt1* “haploid-only” embryos, (E-H) *dseGFP* “haploid-only” embryos, (I-L) *dsDnmt1* “haploid and diploid” embryos, and (M-P) *dseGFP* “haploid and diploid” embryos.

Both ploidy and *Dnmt1* knockdown affected nuclei size during pre-blastoderm development (Fig. 3). Initially, ploidy had an effect on nuclei size with “haploid or diploid” embryos from *dseGFP*-fed females being significantly larger than “haploid-only” embryos from *dseGFP*-fed females (F = 6.0401, p = 0.0142). The “haploid or diploid” embryos from *dseGFP*-fed females had an average nucleus diameter of 10.57 μm (± 0.22 μm) while “haploid-only” embryos from *dseGFP*-fed females had an average nucleus diameter of 8.84µm (± 0.25 µm). Knockdown of *Dnmt1* also significantly reduced nuclei size (F = 162.384, p < 0.001.) Loss of *Dnmt1* resulted in consistently smaller nuclei in both *dsDnmt1* embryo groups (5.98 µm ± 0.335 µm for “haploid-only” embryos and 5.75 µm ± 0.374 µm for “haploid or diploid” embryos). Additionally, there was a significant interaction between ploidy and knockdown of *Dnmt1* (F = 10.436, p = 0.001) with the amplitude of the change in size larger control haploid nuclei due to their larger size in control embryos.

**Fig. 3.**
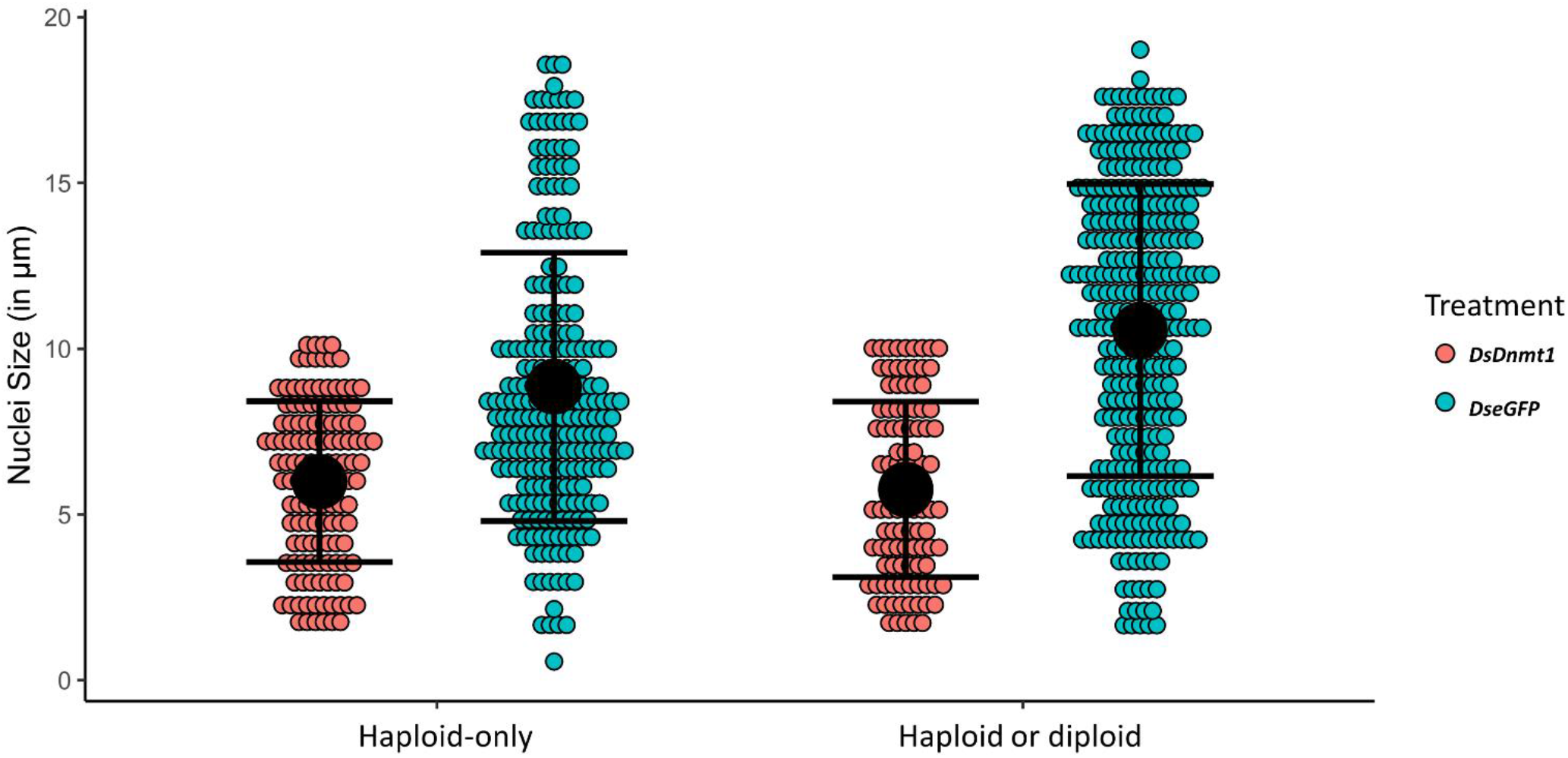
Loss of *Dnmt1* resulted in a smaller nuclei size in both “haploid-only” and “haploid or diploid” embryos. The values are represented as mean ± SE (black circle and bars, respectively) and as individual values (colored circles).

## 4. Discussion

### 4.1. Comparison to embryogenesis in other insects

In this study, we catalogued embryonic development in *B. tabaci*, an emerging model organism for which there are no detailed embryological studies. Based on our descriptions of the timing and morphogenesis during *B. tabaci* embryogenesis, our results indicate that their development is comparable to other insect systems such as *Oncopeltus fasciatus* (Liu & Kaufman, 2004; Panfilio et al., 2006), *Rhodnius prolixus* (Berni et al., 2014), *Gryllus bimaculatus* (Donoughe & Extavour, 2016), *Murgantia histrionica* (Hernandez, Pick, & Reding, 2020), *Nilaparvata lugens* (Fan et al., 2020). This suggests the pattern of development may be conserved amongst hemipterans and perhaps hemimetabolous insects in general. Our study also showed that despite having different nuclei sizes (and presumably, different amounts of DNA), haploid and diploid embryos experienced the same developmental timing, suggesting that ploidy does not control timing in early development alone. This result aligns with what is observed in the hymenopteran *Nasonia*, another obligate haplodiploid system but one that is holometabolous (Arsala & Lynch, 2017). Given their vastly different taxonomic positions, this suggests that haplodiploid insects may rely more heavily on factors other than DNA amount, such as maternal factors, to regulate early development and activate the zygotic genome at the appropriate time to maintain genome function and stability. However, given that maternal factors are not well characterized in this group of insects, it is unclear which major processes (cellularization, cell-cycle pausing, etc.) are being affected in order to coordinate development between haploid and diploid embryos. Overall, embryonic development of *B. tabaci* shares characteristics with other hemimetabolous insects and with other obligate haplodiploid systems.

### 4.2. Comparison to other Dnmt1 knockdown phenotypes

Knockdown of *Dnmt1* prevented *B. tabaci* embryos from completing cellularization and forming a germ rudiment. Cleavage could also have been affected as loss of *Dnmt1* resulted in smaller nuclei. In other insect taxa, loss of *Dnmt1* has resulted in similar embryo failure during the blastoderm phase or prior to gastrulation (Schulz et al., 2018; Ventós-Alfonso et al., 2020; Arsala et al., 2022). This suggests that *Dnmt1* plays an evolutionarily conserved role specifically in early embryogenesis in insects. Indeed, expression of *Dnmt1* peaks during early embryogenesis prior to gastrulation and decreases as development progresses (Arsala et al., 2022). Also, given that loss of *Dnmt1* affected both haploid and diploid embryos prior to the blastoderm phase, it is likely that DNMT1 may be regulating development similar to a maternal factor. The timing and phenotype of the developmental failure could indicate that these embryos are not capable of completing either the transition from maternal to zygotic transcription or the mid-blastula transition, a stage in development characterized by changes in the cell cycle and loss of synchronous cell divisions (Vastenhouw et al., 2019).

### 4.3. Dnmt1 functional considerations

Though *Dnmt1’*s mechanism remains elusive, insight into *when* loss of *Dnmt1* affects development hints at what may be going wrong. The blastoderm phase in most insects is defined by cellularization, zygotic genome activation, and germ rudiment formation. Indeed, these processes are not mutually exclusive as *Drosophila* embryos require zygotic transcripts for cellularization to occur (Edgar et al., 1986) and germ rudiment formation involves differentiation of cell populations in the egg (Johannsen & Butt, 1941). These processes all involve changes in cell cycle regulation. For example, during the cleavage stage, the cell cycle lacks gap phases and rapidly oscillates between DNA synthesis and mitotic divisions (Farrell & O’Farrell, 2014). This occurs because the supply of maternal factors loaded during oogenesis guides the process efficiently without the use of cell cycle checkpoints or much input from the zygotic genome (Reviewed in Brantley & Talia, 2021). The addition of gap phases to the cell cycle coincides with the activation of the zygotic genome and the establishment of cell fate specification domains (Reviewed in Brantley & Talia, 2021). Loss of *Dnmt1* results in downregulation of *Cdc20*, an inducer of mitosis, but not *Cdc25*, which functions during the G2/M checkpoint, suggesting that *Dnmt1* may not be associated with gap phases function, and therefore may not be necessary after the maternal to zygotic transition (Shelby et al., 2023). Indeed, our *Dnmt1* knockdown embryos phenocopy embryos that have lost components involved in maternal to zygotic transition and proper cell cycle progression. For example, loss of function of maternal-to-zygotic transition regulators *Zelda* and *Smaug* also result in failure to cellularize and prevents gastrulation (Arsala & Lynch, 2017). Also, loss of *Piwi* results in mitotic defects that result in abnormal nuclear morphology (Mani et al., 2014) and chromosome condensation and defects during S-phase progression (Schwager et al., 2015). Based on how loss of *Dnmt1* function results in embryo failure during the blastoderm phase and the associated phenotypes, it is likely that plays a role in cell cycle regulation during the maternal to zygotic transition.

Disparate lines of evidence also suggest that DNMT1 helps regulate the cell cycle during embryogenesis in a methylation-independent manner. Studies in mammals have suggested that DNMT1 is involved in related processes such as cell cycle checkpoint recognition and DNA repair (Brown & Robertson, 2007). However, those roles could not be confirmed based on mammalian reliance on a methylated genome, and any resulting phenotype could be attributed to loss of methylation. Loss of *Dnmt1* resulted in loss of reproduction and affected genes related to cell cycle checkpoints even under conditions where not enough mitotic divisions have occurred to reduce DNA methylation (Shelby et al., 2023), suggesting that DNMT1 can interact with DNA without affecting methylation. In our current study, we observed that the syncytial nuclei of knockdown embryos were all small as opposed to being varying sizes based on where they were in the cell cycle at the time of fixation. If nuclei size is suggestive of the cell cycle phase, then our data suggest that the embryos failed at the same point in the cell cycle. However, due to this nuclear phenotype not being reported in other knockdown studies, it is unclear if this phenotype is a result of loss of *Dnmt1* or a random occurrence. Nevertheless, *Dnmt1*’s role in cell cycle activities and genome stability needs to be resolved.

### 4.4. Conclusions and perspectives

In this study, we established a developmental staging system for embryogenesis in *B. tabaci* in order to facilitate comparative and developmental studies. We show that development in *B. tabaci* progresses similarly to that of other hemipterans. We also show that, like other haplodiploid systems, ploidy does not affect developmental timing in *B. tabaci*. Functional assays revealed that *Dnmt1* plays a role in early embryogenesis, suggesting that it may be a maternal regulator of development. In addition, embryos produced from *dsDnmt1*-treated females failed to form a blastoderm and germ rudiment. Because these knockdown embryos had consistent nuclear phenotypes, we suggest that *Dnmt1* may be playing a role in cell cycle checkpoints. However, future studies will need to further study how Dnmt1 interacts with DNA during the cell cycle in order to understand its mode of action.

## Funding

This work was funded by the USDA-ARS Non-Assistance Cooperative Agreement “Managing whiteflies and whitefly-transmitted viruses in vegetable crops in the southeastern U.S.” no. 58-6080-9-006 (P.J.M.). E.A.S. was partially supported by a USDA NIFA National Needs Fellowship award. The mention of a proprietary product does not constitute an endorsement or a recommendation for its use by the USDA or University of Georgia.

## CRediT authorship contribution statement

**Emily A. Shelby:** Writing – original draft, Conceptualization, Methodology, Formal analysis, Data Curation, Investigation, Visualization. **Elizabeth C. McKinney:** Writing – review & editing, Methodology, Validation, Investigation. **Alvin M. Simmons:** Writing – review & editing, Funding acquisition. **Allen J. Moore:** Writing – review & editing, Methodology, Conceptualization, Funding acquisition. **Patricia J. Moore:** Writing-review & editing, Methodology, Supervision, Project administration, Investigation, Conceptualization, Funding acquisition.

## Declaration of competing interest

The authors declare that they have no competing interests.

## Acknowledgements

We thank Dr. Jun-Bo Luan at Shenyang Agricultural University for the whitefly embryo dechorionation protocol. We thank Mr. Charles Dawe for maintaining our whitefly cultures and for growing the plants used in our experiments. We also thank Muthugapatti Kandasamy at the University of Georgia Biomedical Microscopy Core for help with the confocal microscopy.

## Notes

### Competing Interest Statement

The authors have declared no competing interest.

